# GDNF regenerates the missing enteric nervous system of Hirschsprung mice via non-canonical signaling in diverse subtypes of tissue-resident progenitors

**DOI:** 10.1101/2025.05.23.655388

**Authors:** Alassane Gary, Rodolphe Soret, Marie A. Lefèvre, Nejia Lassoued, Ann Aspirot, Christophe Faure, Nicolas Pilon

## Abstract

Hirschsprung disease (HSCR) is a deadly congenital disorder where the enteric nervous system (ENS) is absent from the distal bowel. Current surgical treatment is generally life-saving but is often accompanied by long-term bowel problems and comorbidities. As alternative, we are developing a regenerative therapy based on rectal administration of Glial cell-derived neurotrophic factor (GDNF). We previously showed that administering GDNF enemas to HSCR mice soon after birth is sufficient to permanently induce a new ENS from tissue-resident neural progenitors. Here, we elucidate the underlying mechanism using single-cell transcriptomics, signal transduction inhibitors and genetic cell lineage tracing tools. We found that the neurogenic effect of GDNF is mediated by NCAM1 (Neural cell adhesion molecule 1), rather than by its canonical signaling receptor RET (Rearranged during transfection). We also unveiled the existence of multiple neuronal differentiation pathways that involve a larger than expected repertoire of tissue-resident neural progenitors, including a surprising one not derived from the usual neural crest. These data support feasibility of GDNF-based therapy in most human patients, even those bearing a *RET* variant. This work also has far-reaching implications for the choice of ENS progenitor source to use when developing cell transplantation-based therapeutic approaches.

## INTRODUCTION

The enteric nervous system (ENS) controls motility and other gastrointestinal functions via its vast networks of interconnected neural ganglia ^1–3^. Enteric neurons and glia are both generated from neural crest cell (NCC)-derived progenitors that colonize the developing bowel ^4, 5^. Most progenitors originate directly from vagal NCCs, which enter the foregut mesenchyme and then migrate toward the hindgut to eventually colonize the entire gastrointestinal tract before birth ^6, 7^. When this process fails, ENS ganglia are lacking in the rectum and over varying lengths of the colon, a severe birth defect known as Hirschsprung disease (HSCR) ^5, 8, 9^. NCC-derived Schwann cell precursors (SCPs) constitute another source of ENS progenitors, which in this case use extrinsic nerves as entry points to colonize the bowel, mainly in colon, and after birth ^10, 11^. However, although extrinsic innervation of aganglionic colon in HSCR is more abundant than normal ^12^, the accompanying increase in SCPs cannot spontaneously compensate for the lack of vagal NCC-derived pool ^13, 14^.

Current HSCR treatment still involves surgical removal of the bowel segment lacking ENS ganglia (aganglionic) ^15, 16^. This procedure is generally life-saving but postoperative problems are common ^15, 17–19^. As alternative to this imperfect treatment, we are developing a non-surgical approach based on post-natal stimulation of tissue-resident ENS progenitors ^20, 21^. Using mouse models, we demonstrated that daily rectal administration of GDNF (glial cell line-derived neurotrophic factor) for only 5 days, from post-natal day (P)4 to P8, can regenerate the missing ENS in the colon, with long-term benefits on overall colon function and mouse survival ^20^. Moreover, GDNF promoted *de novo* neurogenesis in human colonic explants obtained from HSCR surgery and cultured *in vitro* ^20^. Our preliminary mechanistic studies suggested that the beneficial effects of GDNF are at least in part due to its ability to promote neuronal differentiation of SCPs from extrinsic nerves ^20^.

During normal pre-natal ENS development, GDNF greatly influences the migration, proliferation and neuronal differentiation of vagal NCCs via the transmembrane tyrosine kinase RET (Rearranged during transfection), upon binding to co-receptor GFRα1 (GDNF family receptor, alpha 1) ^22–25^. This is reflected in HSCR genetics, where *RET* variants constitute the main risk factor of this multifactorial disease ^8, 9^, raising reasonable doubts about global applicability of GDNF-based therapy. Yet, in the Schwann cell lineage, GDNF-bound GFRα1 instead signals through NCAM1 (Neural cell adhesion molecule 1) ^26, 27^, suggesting that RET could in fact be dispensable for mediating postnatal ENS-inducing effects of GDNF in a HSCR context. Clarifying this point will be key to advance GDNF-based therapy closer to clinical application.

A better understanding of cellular heterogeneity among tissue-resident ENS progenitors is also warranted. The majority of GDNF-induced neurons are not derived from SCPs ^20^. Identifying these non-SCP ENS progenitors would allow to better evaluate the requirement for RET signaling, and perhaps also explain how the GDNF-induced ENS is properly balanced in terms of neuron subtypes ^20^ despite that SCP progeny is normally biased toward a cholinergic phenotype ^10^.

## RESULTS

### GDNF changes the proportions of neural cell subpopulations in the distal colon of *Hol^Tg/Tg^* mice

To better understand the mechanism of GDNF-induced neurogenesis in the otherwise aganglionic colon of HSCR mice, we turned to single-cell RNA sequencing (scRNA-seq). We continued to focus on the *Holstein* mouse model for trisomy 21 (collagen VI upregulation)-associated HSCR ^28–30^. This model outperforms two other models available in our laboratory in terms of either phenotypic penetrance of megacolon (compared to incomplete male-biased penetrance in *TashT* line ^31, 32^) or fertility rate (compared to subfertility in *Piebald-Lethal* line ^33^) – both of which being also responsive to GDNF treatment ^20^. To facilitate ENS cell recovery, we bred *Holstein* mice with *G4-RFP* transgenics ^34^. The *G4-RFP* reporter is known to label most NCC derivatives with the red fluorescent protein (RFP) DsRed2 during pre-natal development, including ENS progenitors ^29, 34, 35^. In the most distal quarter of the colon from P10 *Hol^Tg/Tg^;G4-RFP* double transgenic mice that were previously treated or not with GDNF enemas between P4-P8 (Fig.1), we validated that RFP marks with variable intensity 88-90% of SOX10+ glial cells (including extrinsic nerves-associated SCPs) in a GDNF-independent manner (Fig.1a,b) and 70% of GDNF-induced HuC/D+ enteric neurons (Fig.1b) – which are otherwise absent in untreated controls (distal aganglionosis covers ∼25% of colon length in the *Holstein* line ^29^). Of note, unless otherwise indicated, all immunofluorescence stainings in the current study were performed on samples from the last quarter of the colon. Although the percentage of marked neurons was slightly lower than we were hoping, it is noteworthy that similar coverage (73%) did allow to profile a wide variety of postnatal enteric neurons in a recent scRNA-seq study ^36^. Nonetheless, to maximize their recovery by fluorescence-activated cell sorting (FACS), RFP+ cells were sorted in “yield” mode (allowing co-sorting of RFP-negative cells that are close to RFP+ cells in the flow stream) using a non-stringent gating strategy that also included cells with very low levels of RFP signal (in the range of background autofluorescence). These FACS parameters were applied to dissociated cell suspensions from the distal third of the colon (including part of the hypoganglionic transition zone) of 6 untreated control and 8 GDNF-treated mouse pups, from which recovered cells corresponded to ∼1% and ∼2% of all single live cells, respectively. Individual cells were then processed using the 10x Genomics platform (Fig.2a), and subsequently analyzed with the Trailmaker software. This first allowed to separate recovered cells into 19 cell clusters of variable size, including seven associated with the ENS: three with an SCP identity (expressing specific markers like *Dhh, Mpz, Mbp* and *Egfl8*), two with an enteric glial cell (EGC) identity (expressing specific markers like *Nell2, Col20a1, Ccn2* and *Cpe*), and another two with an enteric neuron identity (expressing general markers *Elavl4, Tubb3,* and *Gap43*). Consistent with our non-stringent gating strategy for FACS-mediated recovery, the other cell clusters were not directly associated with the ENS, including specific groups of lymphoid, myeloid, vascular, epithelial and smooth muscle cells. Therefore, to refocus on the ENS, we only selected cells that express either the pan-neuronal marker *Elavl4* or the pan-glial and ENS progenitor marker *Sox10* (Fig.2a), leaving 1354 and 3006 high-quality cells from untreated and GDNF-treated *Hol^Tg/Tg^;G4-RFP* mice, respectively. Unsupervised reclassification of these ENS-associated cells revealed 14 distinct clusters representing subpopulations of SCPs (three clusters, including one with mixed EGC identity), EGCs (eight clusters, including three proliferating) and enteric neurons (three clusters) (Fig.2a-d), which are present in both untreated and GDNF-treated groups (Fig.2e). Analysis of relative proportions revealed a marked enrichment in enteric neurons and proliferating *Mki67*^+^ EGCs in the GDNF-treated group, which occurred mainly at the expense of SCPs and cells with mixed SCP/EGC identity (Fig.2f). These observations are consistent with our prior discovery that GDNF-induced neurons are at least in part derived from SCPs ^20^, thereby making this dataset a reliable resource to better understand the underlying mechanism.

**Figure 1.**
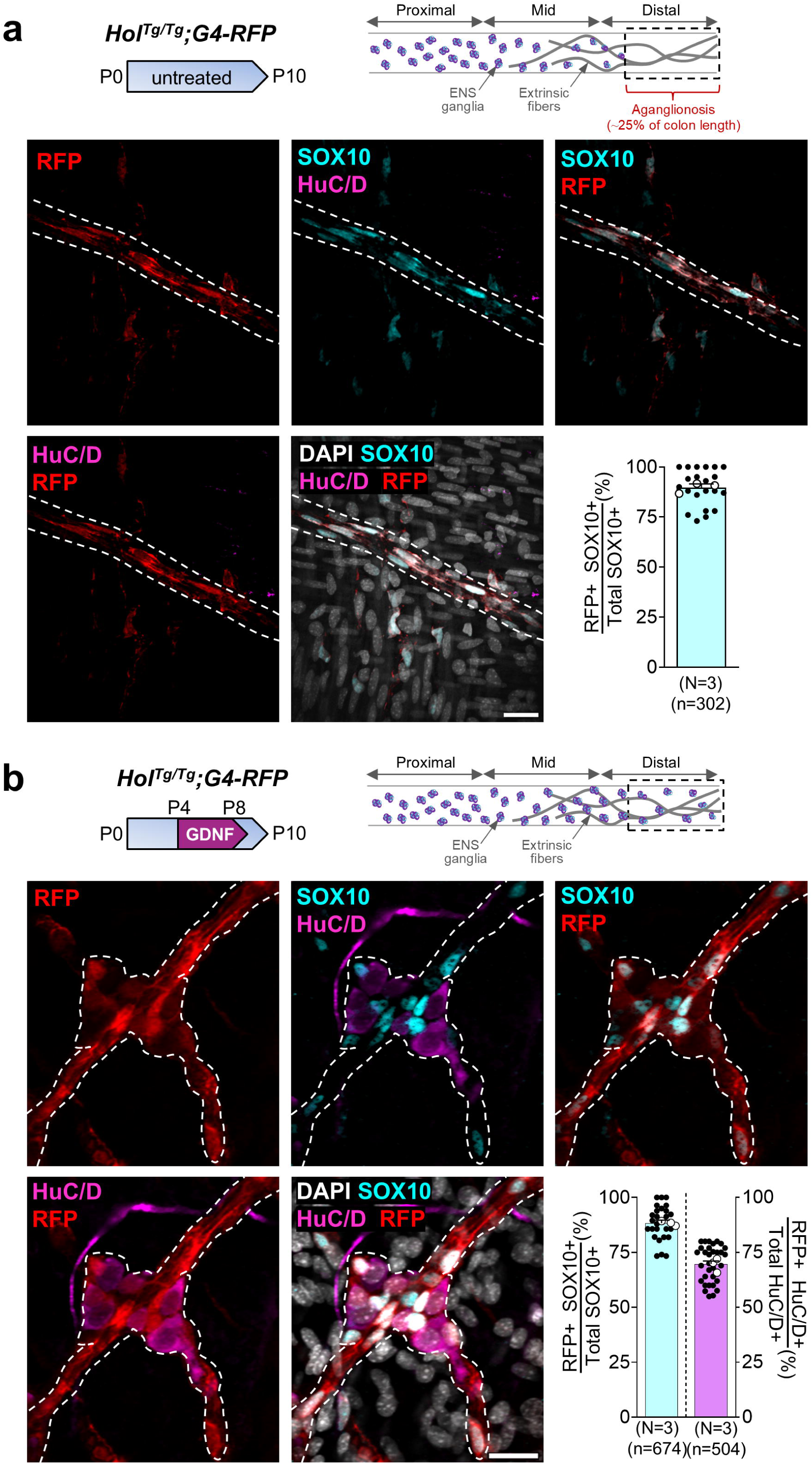
The *G4-RFP* transgene is active in both neurons and glia of the distal colon after birth. **a,b**. Immunofluorescence staining of SOX10, HuC/D and RFP in the distal colon of untreated (a) and GDNF-treated (b) *Hol^Tg/Tg^*;*G4-RFP* at P10. Each black dot in the accompanying quantitative analysis corresponds to a microscopic field of view (7 to 10 fields of view per animal; white dots indicate the average per animal; N indicates the number of animals; n indicates the total number of counted cells). The dashed box in each diagram indicates the colon segment that was analyzed. The dashed outlines in the micrographs highlight extrinsic nerve fibers and/or induced ganglia. Scale bar, 25 μm.

**Figure 2.**
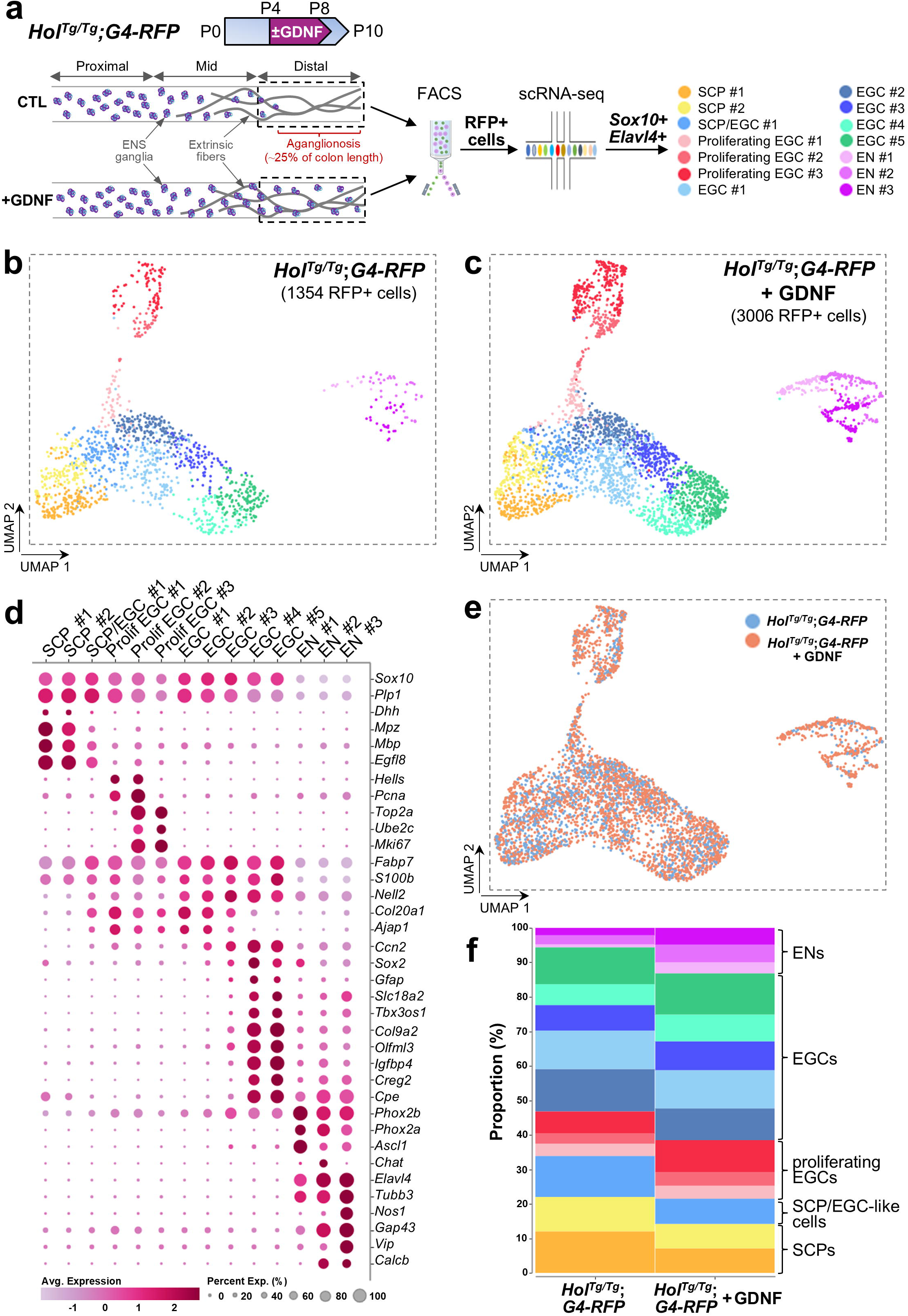
The distal colon of *Hol^Tg/Tg^;G4-RFP* mice is enriched in SCPs and EGCs. **a.** Overview of experimental procedure, from GDNF treatment regimen to scRNA-seq analysis of FACS-recovered RFP+ cells at P10. The dashed boxes in the diagram indicate the colon segment that was analyzed, here also including part of the transition zone. **b,c.** Uniform manifold approximation and projection (UMAP) visualization showing unsupervised clustering of SCPs, EGCs and enteric neurons from distal colon of P10 *Hol^Tg/Tg^;G4-RFP* mice that were treated (c) or not (b) with GDNF. **d.** Dot plot showing relative expression levels of a selection of marker genes in each cluster. **e.** UMAP visualization showing all clusters stratified by experimental group (untreated *vs*. GDNF-treated *Hol^Tg/Tg^;G4-RFP* mice). **f.** Comparative frequency plots showing relative proportion of each cluster in untreated *vs*. GDNF-treated *Hol^Tg/Tg^;G4-RFP* mice.

### The neurogenic effect of GDNF in distal colon is mediated by NCAM1/FAK signaling

We first took advantage of our scRNA-seq data to examine the expression profile of GDNF receptor-coding genes, focusing on GDNF-treated *Hol^Tg/Tg^;G4-RFP* dataset. A striking difference was noted for signaling receptors, with *Ncam1* highly expressed in all cell clusters, while *Ret* expression is mostly enriched in a subset of enteric neurons only (Fig.3a). Binding receptor-coding genes *Gfra1* and *Gfra2* were also found to be widely expressed, with a partly overlapping and complementary distribution (Fig.3a). *Gfra3* expression appeared restricted to SCP clusters, whereas weak *Gfra4* expression could only be detected in a few neurons (Fig.3a). These observations strongly suggested that GDNF could bind either GFRα1 or GFRα2 ^37^ and then signals through NCAM1 to make new neurons from SCPs and perhaps other ENS progenitors within the EGC clusters. To further validate this possibility, we evaluated the distribution of the NCAM1 protein and its downstream signaling effector phospho-FAK[Y397] in most distal colon of *Hol^Tg/Tg^* mice over time, before (P4_[0h]_), during (P4_[6h]_ and P6) and after (P10) the end of GDNF treatment. NCAM1 is already present in SOX10+ cells before GDNF treatment begins at P4, both within and outside extrinsic nerve fibers (Fig.3b,c). As soon as 6h after first GDNF administration, NCAM1 levels then increase over time, with high levels being maintained even two days after completion of treatment period (Fig.3b,c). Phospho-FAK[Y397] levels are lower in these SOX10+ cells before the start of GDNF treatment, but then similarly increase over time (Fig.3b,c). These changes in NCAM1 and phospho-FAK[Y397] levels do not appear to be due to a simple temporal effect, as we did not observe overt changes in the ENS-containing mid-colon of untreated *Hol^Tg/Tg^* mice between P4 and P10 (Fig.3d). Hence, these additional observations suggest that NCAM1 could also be subjected to GDNF-induced autoregulation, as we previously showed for RET ^20^.

**Figure 3.**
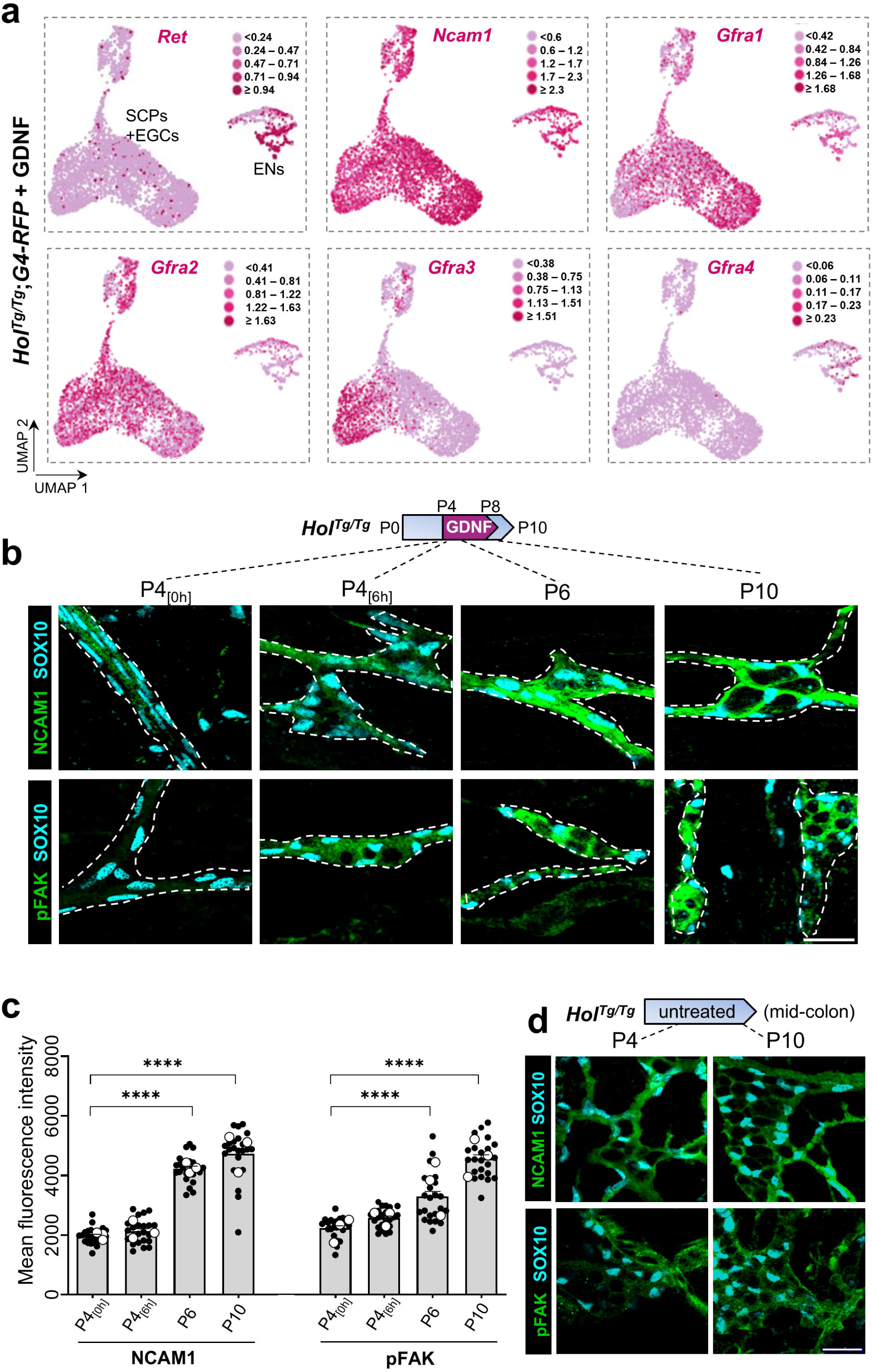
NCAM1 and phospho-FAK[Y397] are stimulated by GDNF in the distal colon of *Hol^Tg/Tg^;G4-RFP* mice. **a.** Feature plots showing relative gene expression levels of signaling (*Ret* and *Ncam1*) and binding (*Gfra1, Gfra2, Gfra3, Gfra4*) GDNF receptors in scRNA-seq dataset of GDNF-treated *Hol^Tg/Tg^;G4-RFP* mice. **b,c.** Immunofluorescence staining of NCAM1 and its downstream signaling effector phospho-FAK[Y397] in most distal colon of *Hol^Tg/Tg^* mice at indicated time points before, during and after GDNF treatment (panel b shows representative images of N=3 animals per time point). Quantification of mean fluorescence intensity for each experimental group is shown in panel c. Each black dot corresponds to a single microscopic field of view (7–8 imaging fields per animal; white dots indicate the average per animal). \*\*\*\**P* < 0.0001; Statistics were calculated from the datasets plotted as black dots using ordinary one-way ANOVA with post-hoc Tukey’s test. Dashed outlines in panel b highlight either extrinsic nerve fibers or individual ganglia. **d.** Immunofluorescence staining of NCAM1 and phospho-FAK[Y397] in mid-colon of untreated *Hol^Tg/Tg^* mice at indicated time points (representative images of N=3 animals per time point). Scale bar, 25 μm.

To confirm the relative importance of NCAM1 *vs.* RET in postnatal colon of *Hol^Tg/Tg^* mice, we next verified GDNF treatment efficacy in presence of chemical inhibitors that prevent phosphorylation of FAK (PF-562271)^38^ and RET (BLU-667)^39^, respectively. Each of these inhibitors was administered via daily i.p. injections (30 mg/kg) during the GDNF enema treatment period (P4-P8), and impact on colon tissues was then assessed 6h after last enema/injection at P8 (Fig.4a). Western blot analysis of full-thickness proximal/mid colon confirmed efficacy of each inhibitor (Fig.4b), while further highlighting that RET is poorly activated in this context compared to a control tissue with robust endogenous GDNF-RET activity like the hypothalamus ^40^ (Fig.4b). Accordingly, immunofluorescence staining of most distal colon segment from same animals revealed that phospho-RET[Y1062] inhibition with BLU-667 has no impact on GDNF-induced neurogenesis. In marked contrast, phospho-FAK[Y397] inhibition with PF-562271 led to a significant decrease in the number of GDNF-induced neurons per mm² (Fig.4c), without inducing apoptosis in the ENS (Fig.4d). Rare apoptotic neurons positive for cleaved Caspase 3 were only detected in presence of the phospho-RET[Y1062] inhibitor BLU-667 (Fig.4d). These data confirm that NCAM1/FAK signaling is crucial for GDNF-induced neurogenesis in aganglionic colon of a HSCR mouse model.

**Figure 4.**
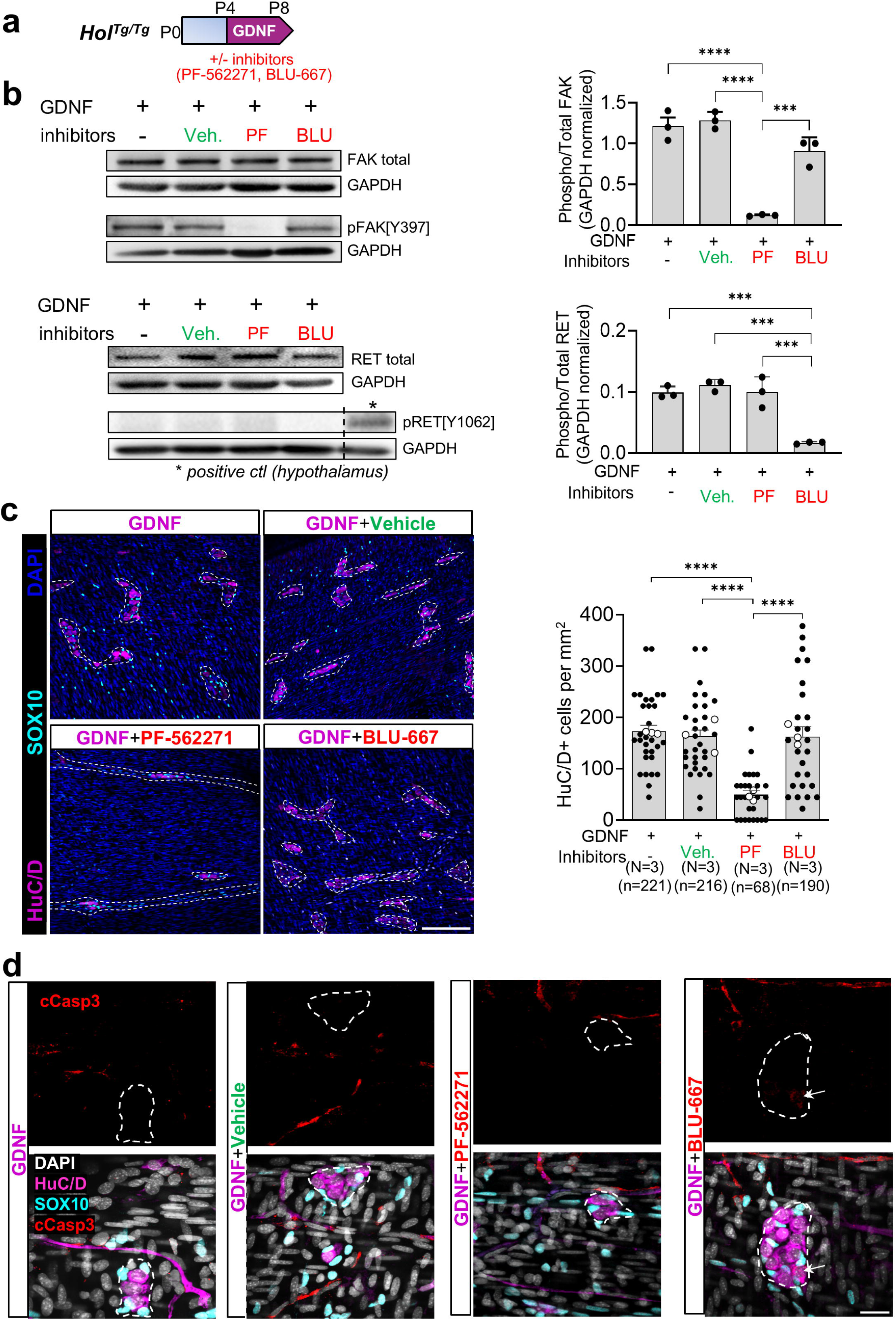
The neurogenic effect of GDNF in *Hol^Tg/Tg^;G4-RFP* mice is mediated by NCAM1, not RET. **a.** Diagram of the experimental procedure showing that *Hol^Tg/Tg^* mice were administered GDNF enemas and were also injected with the indicated selective inhibitors (or vehicle only) for phospho-FAK[Y397] (PF-562271) or phospho-RET[Y1062] (BLU-667) from P4 to P8. **b.** Western blot analysis of both total and phosphorylated forms of FAK or RET in the proximal colon of P8 *Hol^Tg/Tg^* mice treated as indicated above. Each value in the accompanying quantitative analyses of phospho/total ratios correspond to an animal. \*\*\**P* < 0.001; \*\*\*\**P* < 0.0001; Statistics were calculated from the datasets plotted as black dots using ordinary one-way ANOVA with post-hoc Tukey’s test. **c,d.** Immunofluorescence staining of HuC/D and SOX10 (c) and cleaved Caspase 3 (d) in most distal colon of the same P8 *Hol^Tg/Tg^* mice that were used in panel b. Quantitative analysis of HuC/D+ neurons per mm^2^ is shown on the right, with each black dot corresponding to a microscopic field of view (7 to 10 fields of view per animal; white dots indicate the average per animal; N indicates number of animals per group; n indicates total number of counted cells). \*\*\*\**P* < 0.0001; Statistics were calculated from the datasets plotted as black dots using ordinary one-way ANOVA with post-hoc Tukey’s test. Dashed outlines in panels c-d highlight either extrinsic nerve fibers or individual ganglia. Scale bar, 150 µm (c) and 25 μm (d).

### GDNF-induced neurogenesis from distal colon SCPs partly involves an intermediary EGC-like state

We reasoned that our scRNA-seq data would also be helpful to identify additional sources of GDNF-induced neurons, other than SCPs ^20^. Indeed, as some EGCs behave as ENS progenitors with a neurogenic potential under certain conditions ^14, 41–45^, we hypothesized that at least one of the many subgroups of EGCs in our analysis of P10 mice would contain such progenitors. To uncover neuronal-fated progenitors, we verified the expression pattern of the late pan-neuronal marker *Elavl4*, predicting that it should start to be upregulated while the EGC transcriptional program is still active. Again focusing on the GDNF-treated *Hol^Tg/Tg^;G4-RFP* dataset, this analysis confirmed that several *Elavl4* expressing cells are among both SCP and EGC clusters, in addition to all enteric neuron clusters (Fig.5a). Reclustering these *Elavl4* expressing cells then unveiled an intriguing connection between SCP, EGC and enteric neuron clusters (Fig.5b-d). This linear relationship was corroborated by a pseudotime trajectory analysis (without predicting directionality), which further confirmed it exists in untreated *Hol^Tg/Tg^;G4-RFP* dataset as well (Fig.5e) – most likely reflecting presence of ENS-containing transition zone in our samples. Moreover, we observed a similar linear connection when the reclustering was based on *Tubb3* expression instead of *Elavl4*, with overall inverse expression gradients of *Sox10* and these pan-neuronal markers (Fig.5d). Considering all of the above, these analyses suggest the existence of a stepwise differentiation model (Fig.5c-d) whereby a subset of SCPs (expressing *Dhh*, *Mpz, Mbp* and *Egfl8*) progressively loose their Schwann cell lineage identity to differentiate into EGCs expressing *Nell2, Col20a1* and *Ajap1* (SCP/EGC and EGC #a clusters), then EGCs expressing *Ccn2*, *Col9a2* and *Igfbp4* (EGC #b cluster), and finally enteric neurons expressing general markers like *Phox2b, Tubb3* and *Gap43,* as well as different combinations of subtype-specific markers like *Chat, Nos1, Vip* and *Calcb* (EN #a-f clusters). Interestingly, the EGC #b cluster also expresses *Gfap*, *Slc18a2, Sox2* and *Cpe* (Fig.5c-d). These genes collectively characterize EGCs poised for neurogenesis in the small intestine of *Plp1-GFP* mice at P14 ^46^. Moreover, these poised EGCs were recently assigned to the “EG 1-2” clusters identified in pediatric human ileum and colon biopsies ^47^.

**Figure 5.**
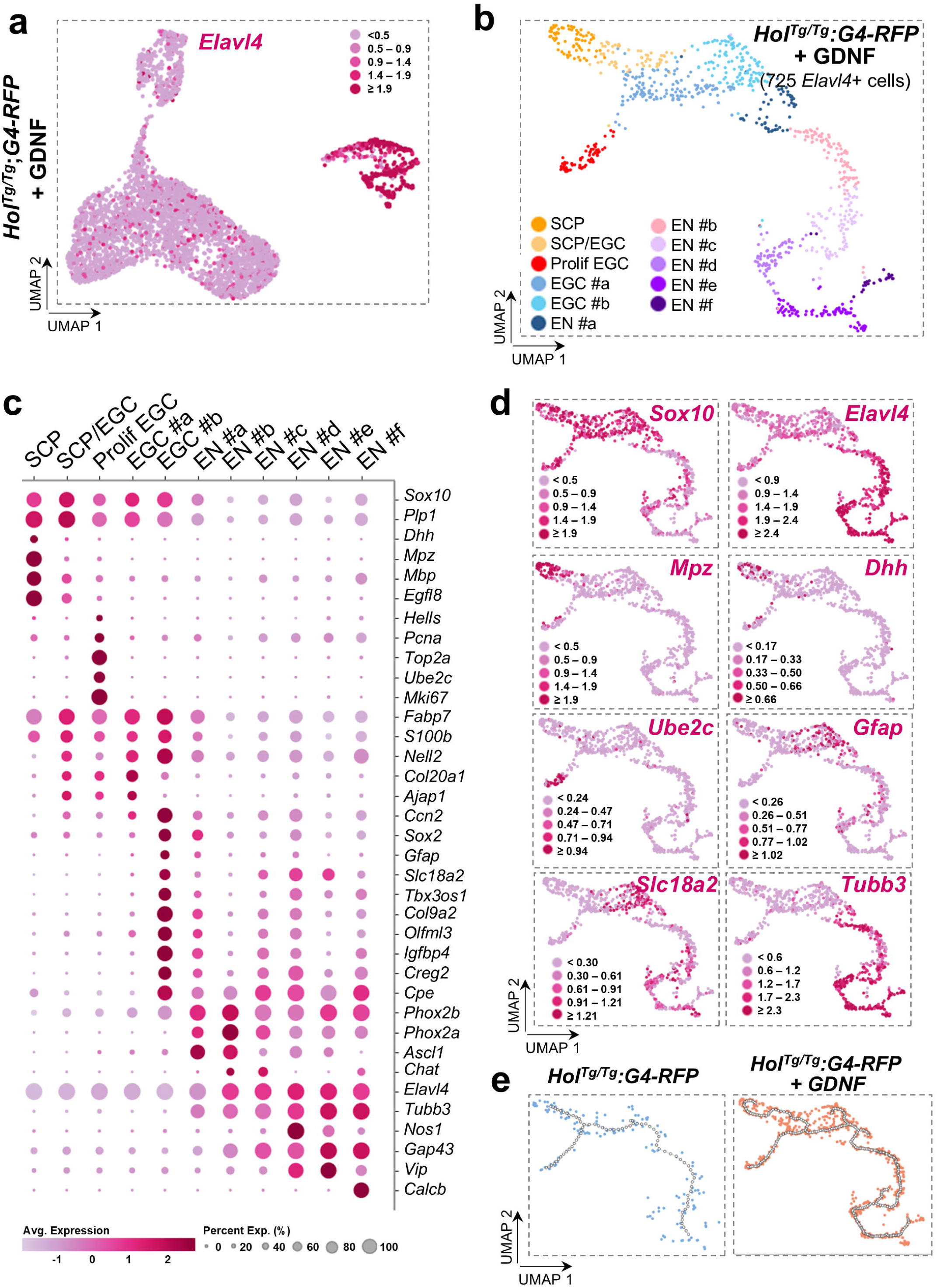
Re-clustering based on *Elavl4* expression reveals a stepwise differentiation mechanism connecting SCPs, EGCs and enteric neurons. **a.** Feature plot showing *Elavl4* expression levels in scRNA-seq dataset of GDNF-treated *Hol^Tg/Tg^;G4-RFP* mice. **b.** UMAP visualization of scRNA-seq dataset of GDNF-treated *Hol^Tg/Tg^;G4-RFP* mice after reclustering based on *Elavl4* expression levels. **c.** Dot plot showing relative expression levels of a selection of marker genes in each cluster. **d.** Feature plots showing relative gene expression levels of pan-glial (*Sox10*), pan-neuronal (*Elavl4* and *Tubb3*), SCP (*Mpz* and *Dhh*), cycling cell (*Ubec2c*) and EGC (*Gfap* and *Slc18a2*) markers. e. Pseudotime analysis of *Elavl4*-based reclustered scRNA-seq datasets of untreated and GDNF-treated *Hol^Tg/Tg^;G4-RFP* mice.

To validate the stepwise differentiation model described above at both genetic and protein levels, we turned again to a cell lineage tracing system that previously allowed us to follow progeny of *Dhh*-expressing SCPs in different contexts ^11, 14, 48, 49^, including *Hol^Tg/Tg^* mice treated with GDNF ^20^. In these GDNF-treated triple-transgenic *Hol^Tg/Tg^* pups also bearing the *Dhh-Cre* transgene ^50^ and the *R26R-YFP* reporter allele ^51^ (*Hol^Tg/Tg^;Dhh-Cre^Tg/+^;R26^YFP/+^*), we now evaluated the presence of SCP-derived cells co-expressing either GFAP or SLC18A2 proteins together with the early enteric neuron marker PHOX2B in the distal colon at P10 (*i.e.* 2 days after last GDNF enema, as for scRNA-seq). Triple-immunofluorescence staining clearly shows that GDNF-induced ENS ganglia contain several SCP-derived (YFP+) cells that are differentiating into PHOX2B+ enteric neurons (as also evidenced by their typical large, rounded nucleus) while still expressing EGC markers GFAP (Fig.6a) or SLC18A2 (Fig.6b). Yet, quantitative analysis as a function of time revealed that GFAP+ cells in the distal colon are not all derived from *Dhh*-expressing SCPs on the first day of treatment (Fig.6c). Furthermore, this kinetic analysis highlighted a progressive decline of GFAP protein expression in SCP-derived cells in the distal colon during (P4_[6h]_ and P6) and after (P10 and P20) GDNF treatment, with a mean of 87% of YFP+ cells per ganglion also positive for GFAP after the first 6h of treatment to a mean of 31% at P20 (Fig.6c). This progressive decline most likely reflects the inverse progression of neuronal differentiation, as GFAP is no longer detectable in SCP-derived cells that stain positive for the late neuronal marker HuC/D at P20 (Fig.6d). Collectively, these results support the notion that some extrinsic nerve-associated SCPs from the distal colon can differentiate into enteric neurons by transitioning through an EGC-like state.

**Figure 6.**
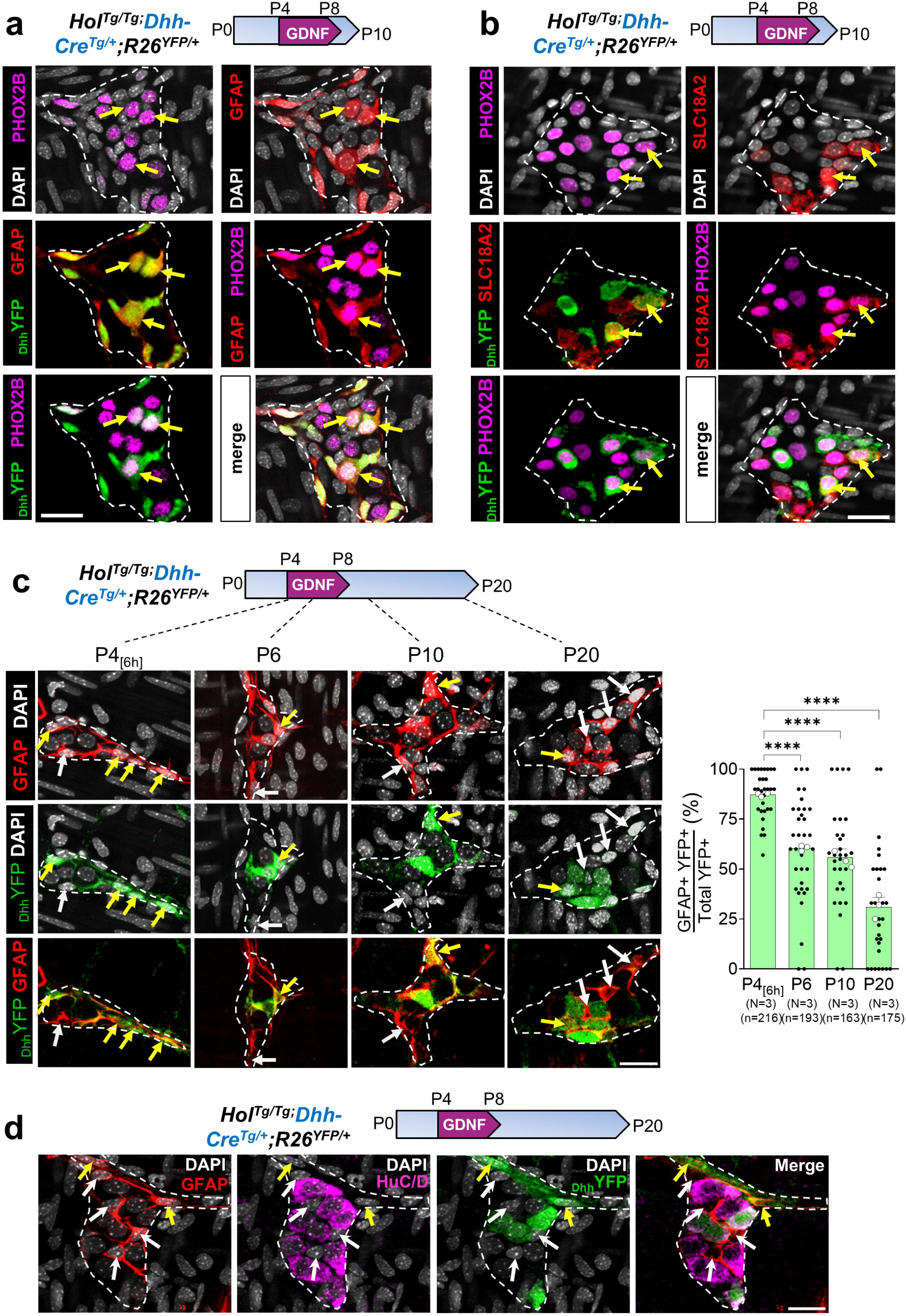
GDNF-induced neuronal differentiation of SCPs occurs in part via and EGC-like state. **a,b.** Immunofluorescence staining of most distal colon samples from P10 GDNF-treated *Hol^Tg/Tg^;Dhh-Cre^Tg/+^;R26^YFP/+^* mice showing several newly induced myenteric ganglion cells derived from SCPs (YFP+) that are undergoing neuronal differentiation (PHOX2B+ cells with large rounded nucleus) while also expressing either GFAP (a) or SLC18A2 (b) EGC markers (see yellow arrows; representative images of N=3 animals). **c,d.** Immunofluorescence staining of most distal colon samples from GDNF-treated *Hol^Tg/Tg;^Dhh-Cre^Tg/+^;R26^YFP/+^* mice at indicated time points showing that proportion of SCP-derived (YFP+) myenteric ganglion cells also positive for the EGC marker GFAP decreases over time (c) and that GFAP is not expressed in mature neurons at P20 (d). Yellow and white arrows point to GFAP+ cells that are either YFP+ or YFP-, respectively. Quantitative analysis of percentage of double YFP+ GFAP+ myenteric ganglion cells is shown on the right of panel c, with each black dot corresponding to a microscopic field of view (7 to 10 fields of view per animal; white dots indicate the average per animal; N indicates number of animals per group; n indicates total number of counted cells). \*\*\*\**P* < 0.0001; Statistics were calculated from the datasets plotted as black dots using ordinary one-way ANOVA with post-hoc Tukey’s test. Dashed outlines in panels a-d highlight individual ganglia. Scale bar, 25 μm.

### GDNF-induced neurogenesis in the distal colon is quicker from SCPs than from EGCs and persists beyond the treatment period

To clarify the relationship between distal colonic SCPs and EGCs in response to GDNF treatment of *Hol^Tg/Tg^* mice, we extended our cell lineage tracing studies by comparing the kinetic profile of GDNF-induced neurogenesis from *Dhh*-expressing SCPs as well as from EGCs expressing either *Gfap* (with tamoxifen-inducible *GFAP-CreERT2* driver) ^52^ or *Slc18a2* (with the *Slc18a2-Cre* driver) ^53^. Like most of our other kinetic analyses, we analyzed one time point just before treatment begins (P4_[0h]_), two time points during the GDNF treatment period (P4_[6h]_ and P6), and two time points after the end of treatment (P10 and P20) (Fig.7a). Regardless of their origin, HuC/D+ neurons in distal colon became detectable by immunofluorescence remarkably quickly, as early as 6h after the start of treatment (Fig.7b). Otherwise undetectable in this most distal colon region without exposure to GDNF, these HuC/D+ neurons began to appear either isolated or organized into very small ganglia (Fig.7c-e). At this very early stage, they were detected in all distal colon samples of GDNF-treated pups, but not in all fields of view (with 60x objective). After this P4_[6h]_ time point, the mean density of HuC/D+ neurons per mm^2^ increased in a steady manner and, surprisingly, even well after the 5-day GDNF treatment period (P4-P8) ended, eventually reaching a plateau from P20 onwards (Fig.7b).

**Figure 7.**
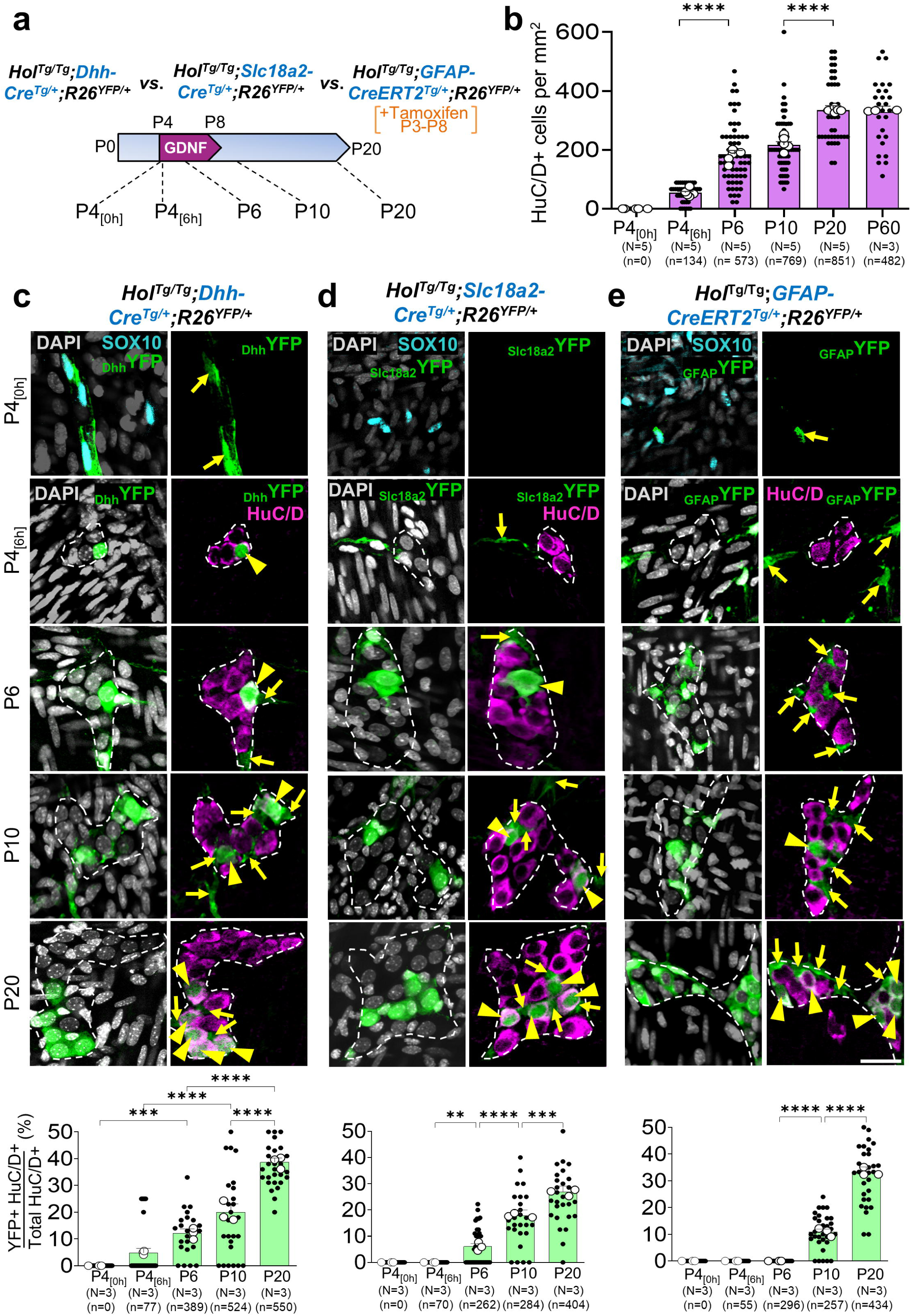
GDNF-induced neurogenesis is more rapid from SCPs than from EGCs. **a.** Diagram of time-course experimental procedure and involved mouse models for comparative genetic cell lineage tracing in the *Hol^Tg/Tg^* mutant background. **b.** Quantitative analysis of density of GDNF-induced myenteric neurons in most distal colon of *Hol^Tg/Tg^* mice as a function of time, regardless of cell lineage tracing tools. **c-e.** Immunofluorescence-based kinetic analysis of distal colon myenteric SOX10+ glia (yellow arrows) or HuC/D+ neurons (yellow arrowheads) that are also positive for YFP, which indicates whether they are derived from *Dhh*-expressing SCPs (c), *Slc18a2*-expressing EGCs (d) or *Gfap*-expressing EGCs (e). For all quantitative analyses, each black dot corresponds to a microscopic field of view (7 to 11 fields of view per animal; white dots indicate the average per animal; N indicates number of animals; n indicates total number of counted cells). \**P* < 0.5; \*\*\**P* < 0.001; \*\*\*\**P* < 0.0001; Statistics were calculated from the datasets plotted as black dots using ordinary one-way ANOVA with post-hoc Tukey’s test. Dashed outlines in panels c-e highlight individual ganglia. Scale bar, 25 μm.

When considering Cre drivers used for cell lineage tracing, our comparative kinetic analysis revealed that *Dhh*-expressing SCPs are among the first newly generated HuC/D+ neurons, after only 6h of treatment (Fig.7c). GDNF-induced neurogenesis became apparent much later in EGCs expressing either *Slc18a2* (Fig.7d) or *Gfap* (Fig.7e), with first detection of YFP+ HuC/D+ neurons at P6 and P10, respectively. Before these stages, both of these Cre drivers allow to label a subset of SOX10+ EGCs with the YFP reporter (Fig.7d,e), but no neurons. On one hand, this temporal difference in neurogenesis thus supports the stepwise differentiation model (SCP → EGC → enteric neurons) described above (Figs.5-6). On the other hand, as *Gfap*-expressing EGCs most likely exist independently of the SCP lineage (Fig.6c), it also suggests that neuronal differentiation in distal colon is simply slower from EGCs than from SCPs. After first detection, percentage of YFP+ neurons then increases over time with all three Cre drivers, in a way roughly similar to the overall time-dependent increase in neuronal density (Fig.7b), with final contribution at P20 reaching between 27-39% of all GDNF-induced neurons (Fig.7c-e). Hence, most neurons originating from SCPs and/or EGCs (at least those labeled with our cell lineage tracing tools) are generated after the GDNF treatment period between P4-P8. During the GDNF treatment period, most of the newly induced neurons instead seem to have another origin, distinct from *Dhh*-expressing SCPs or *Slc18a2/Gfap*-expressing EGCs.

### Direct transdifferentiation is the preferred mode of neurogenesis in response to GDNF, regardless of targeted ENS progenitor subtype

The rapid neuronal differentiation uncovered at P4_[6h]_ (Fig.7b) strongly suggested that the first GDNF-induced neurons that appear in the distal colon of *Hol^Tg/Tg^* mice result from direct transdifferentiation, without cell division. To validate this hypothesis and more globally evaluate the contribution of such direct neuronal conversion, we conducted EdU incorporation assays in GDNF-treated *Hol^Tg/Tg^* pups, again as a function of time and subtypes of ENS progenitors. Daily EdU treatment began at P3 and continued throughout duration of GDNF treatment period (Fig.8a). As expected, immunofluorescence staining of distal colon showed that EdU incorporation into GDNF-induced HuC/D+ neurons was extremely rare at P4_[6h]_ (Fig.8b), in contrast to numerous proliferative cells in their vicinity. We found only two neurons in total, from the same mouse, that stained positive for EdU at this stage. Extending this global analysis at later time points revealed a time-dependent increase in EdU+ neurons, but these always remained the minority, resulting in consistently high percentages (>67%) of direct conversion (*i.e.*, without EdU incorporation), both during (P6) and after (P10 and P20) the GDNF treatment period (Fig.8b). Similar trends can be observed when data are analyzed based on Cre drivers used for lineage tracing (Fig.8c-e). Percentages of EdU-negative neurons in distal colon are always the highest at first detection of YFP+ neurons, with relative levels being inversely proportional to the stage at which this occurs in each mouse line: 100% at P4_[6h]_ for *Dhh*-expressing SCPs (Fig.8c), 91% at P6 for *Sclc18a2*-expressing EGCs (Fig.8d), and 77% at P10 for *Gfap*-expressing EGCs (Fig.8e). These data offer a plausible mechanism to explain the rapid generation of GDNF-induced neurons after just 6h of treatment, while also indicating that direct transdifferentiation remains the preferred mode of neurogenesis irrespective of temporal constraints.

**Figure 8.**
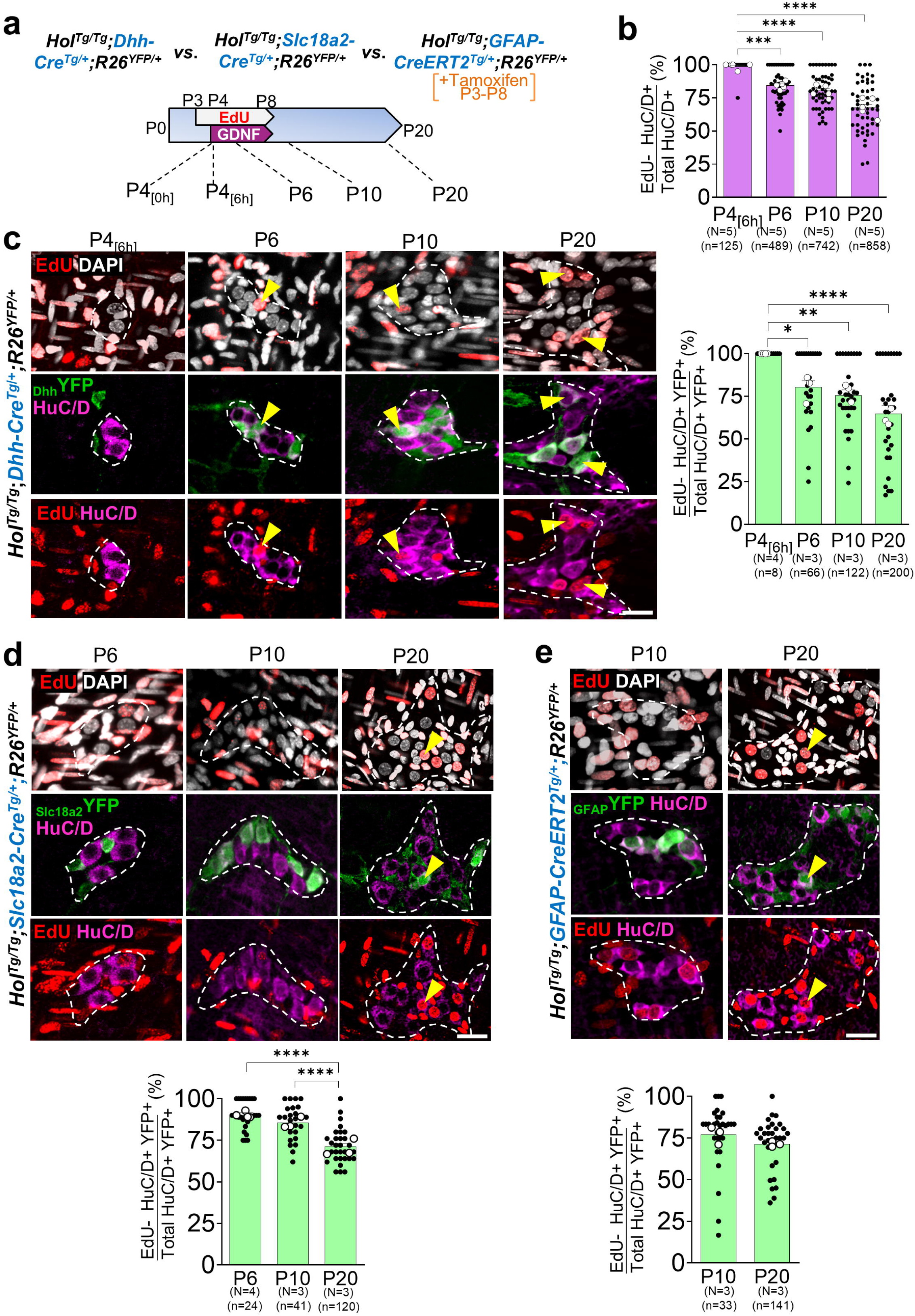
GDNF predominantly promotes direct transdifferentiation of ENS progenitors into neurons. **a.** Diagram of time-course experimental procedure and involved mouse models for combined EdU incorporation assays and genetic cell lineage tracing in *Hol^Tg/Tg^* mutant background. **b.** Quantitative analysis of percentage of GDNF-induced HuC/D+ myenteric neurons that did not incorporate EdU in most distal colon of *Hol^Tg/Tg^* mice as a function of time, regardless of cell lineage tracing tools. **c-e**. Immunofluorescence-based kinetic analysis of distal colon myenteric HuC/D+ neurons (yellow arrowheads) that have incorporated EdU and are also positive for YFP, which indicates whether they are derived from *Dhh*-expressing SCPs (c), *Slc18a2*-expressing EGCs (d) or *Gfap*-expressing EGCs (e). For all quantitative analyses, each black dot corresponds to a microscopic field of view (7 to 11 fields of view per animal; white dots indicate the average per animal; N indicates number of animals; n indicates total number of counted cells). \**P* < 0.5; \*\*\**P* < 0.001; \*\*\*\**P* < 0.0001; Statistics were calculated from the datasets plotted as black dots using ordinary one-way ANOVA with post-hoc Tukey’s test (c, d) or unpaired two-tailed Student’s *t*-test (e). Dashed outlines in panels c-e highlight individual ganglia. Scale bar, 20 μm.

### A neural crest-independent ENS progenitor subtype appears necessary to balance overall cholinergic and nitrergic differentiation of GDNF-induced neurons in the distal colon

Finally, we sought to determine the specific contribution of SCPs and EGCs to WT-like diversity of neuronal subtypes that we know GDNF can generate in the distal colon of *Hol^Tg/Tg^* mice ^20^. Strikingly, our analyses at P20 showed that the vast majority of YFP+ neurons derived from either *Dhh*-expressing SCPs or *Gfap-*expressing EGCs are also positive for the excitatory motor neuron marker ChAT (choline acetyltransferase), with proportions of 83% and 80%, respectively (Fig.9a,b). Much smaller percentages of YFP+ neurons were found to co-express the inhibitory motor neuron marker NOS1 (nitric oxide synthase 1), with averages of 12% for those derived from *Dhh*-expressing SCPs and 14% from *Gfap*-expressing EGCs (Fig.9a,b). Similar trends were noted with the *Slc18a2*-Cre driver (Fig.9c), with a slightly higher percentage of nitrergic neurons that most likely reflects, at least in part, *Slc18a2* expression in a subset of these neurons (Fig.2d). When compared to each other, data for *Dhh*-expressing SCPs and *Gfap-*expressing EGCs result in cholinergic to nitrergic ratios of 6.3 : 1 on average, which is much greater than the WT-like ratio of 1.2 : 1 previously reported for all GDNF-induced HuC/D+ neurons (52% ChAT+ *vs*. 42% NOS1+) ^20^. This ratio difference led us think that the additional subtype of ENS progenitors already predicted from our temporal cell lineage tracing data (Fig.7) must be more prone to differentiate into nitrergic neurons.

**Figure 9.**
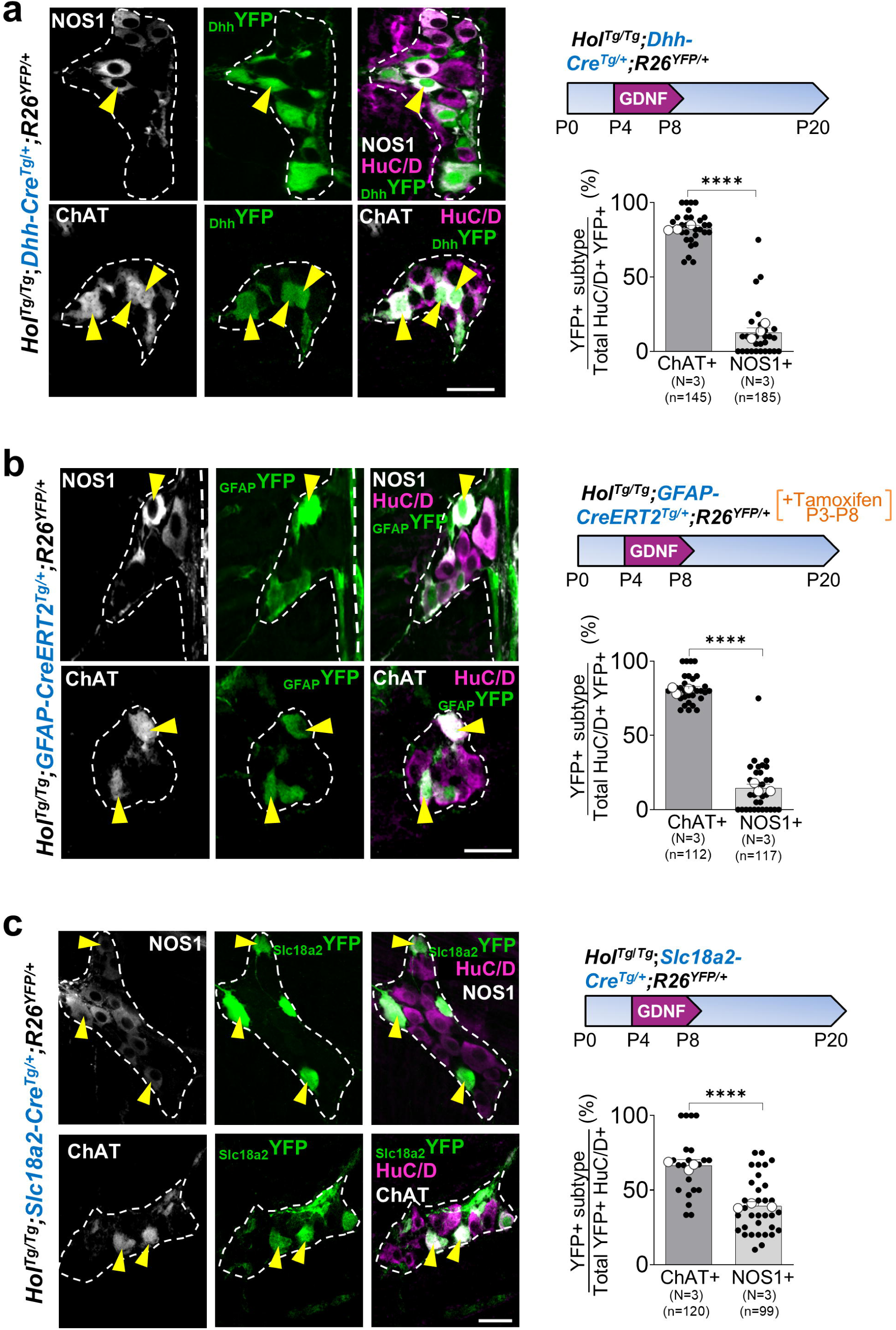
GDNF-induced neurons derived from *Dhh*-expressing SCPs and *Gfap*-expressing EGCs are mostly cholinergic. **a-c.** Immunofluorescence-based analysis of relative proportions of nitrergic (NOS1+) and cholinergic (ChAT+) neurons among GDNF-induced myenteric neurons (HuC/D+) in the most distal colon at P20, with YFP signal (yellow arrowheads) further indicative of their origin from *Dhh*-expressing SCPs (a), *Gfap*-expressing EGCs (b) or *Slc18a2*-expressing cells (c). For all quantitative analyses, each black dot corresponds to a microscopic field of view (7 to 11 fields of view per animal; white dots indicate the average per animal; N indicates number of animals; n indicates total number of counted cells). \*\*\*\**P* < 0.0001; Statistics were calculated from the datasets plotted as black dots using unpaired two-tailed Student’s *t*-test. Dashed outlines highlight individual ganglia. Scale bar, 25 μm.

To verify that this presumed additional progenitor subtype could be derived from NCCs but was simply missed by our scRNA-seq strategy based on *G4-RFP* activity (Figs.1 and 2a), we traced all NCC derivatives in the colon using the *Wnt1-Cre2* driver ^54^. When used in conjunction with the *R26R-YFP* reporter allele (*Wnt1-Cre2^Tg/+^;R26^YFP/+^*), this “pan-NCC” Cre driver normally enables YFP expression in virtually all enteric neurons and EGCs of distal colon at P20, as we confirmed ourselves (Fig.10a). However, this was not the case when the same system was used in the *Holstein* mutant background. To our astonishment, about a quarter of GDNF-induced HuC/D+ neurons were negative for YFP in the distal colon of these *Hol^Tg/Tg^;Dhh-Cre^Tg/+^;R26^YFP/+^* mice at P20 (Fig.10a). This ENS contribution from a non-NCC source appeared specific to the distal colon (not observed in already ganglionated proximal colon of the same mice; see insets in Fig.10a) and was strongly oriented toward neurons: less than 3% of SOX10+ EGCs were similarly negative for YFP in the same GDNF-treated animals (Fig.10a). Importantly, this atypical non-NCC-related subtype of ENS progenitors appeared ∼4X more prone to differentiate into nitrergic neurons, now yielding roughly equal proportions of NOS1+ and ChAT+ neurons (Fig.10b).

**Figure 10.**
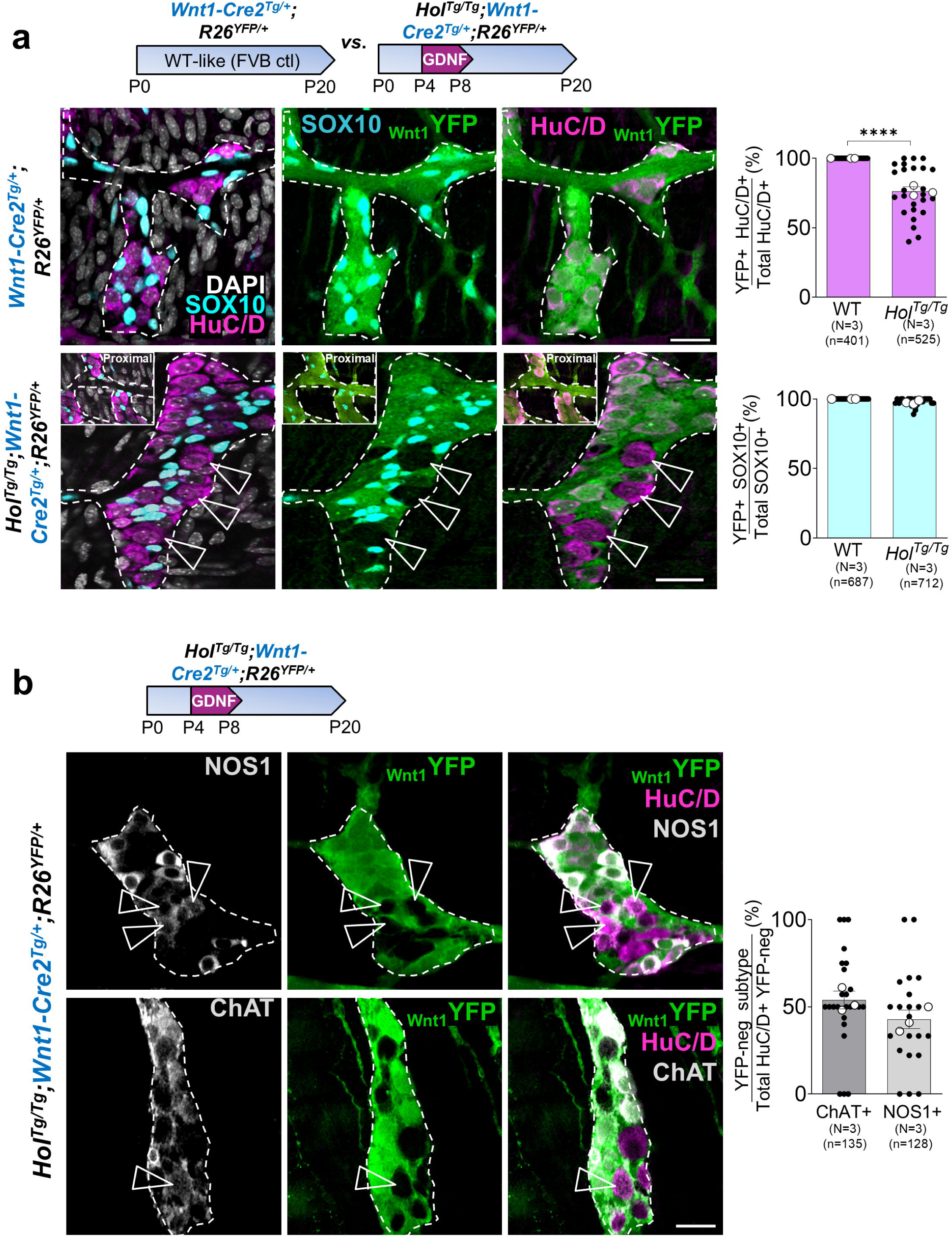
A neural crest-independent ENS progenitor subtype is required for balancing cholinergic *vs*. nitrergic differentiation of GDNF-induced myenteric neurons. **a.** Immunofluorescence-based analysis of the contribution of *Wnt1*-expressing NCCs (YFP+) to overall pool of HuC/D+ neurons and SOX10+ EGCs in most distal colon of P20 WT-like control and GDNF-treated *Hol^Tg/Tg^* mutant mice. Empty arrowheads point to YFP-negative neurons in *Hol^Tg/Tg^* mutant background. Insets show results for the proximal colon of the same mouse as internal positive control **b.** Immunofluorescence-based analysis of relative proportions of nitrergic and cholinergic neurons among GDNF-induced myenteric neurons in the most distal colon at P20, with YFP signal further indicative of their origin from *Wnt1*-expressing NCCs. Empty arrowheads point to YFP-negative neurons. For all quantitative analyses, each black dot corresponds to a microscopic field of view (7 to 11 fields of view per animal; white dots indicate the average per animal; N indicates number of animals; n indicates total number of counted cells). \*\*\*\**P* < 0.0001; Statistics were calculated from the datasets plotted as black dots using unpaired two-tailed Student’s *t*-test. Dashed outlines highlight individual ganglia. Scale bar, 25 μm.

## DISCUSSION

In this study, we greatly deepened our understanding of GDNF-induced neurogenesis in the context of HSCR. Just a few hours after rectal administration, GDNF triggers a robust neurogenic response in the distal colon, which then persists for several days even when GDNF is no longer administered. This GDNF-induced neurogenic program is mediated by NCAM1-FAK signaling, not by RET. Within the aganglionic distal colon, at least three distinct subtypes of ENS progenitors are likely involved (four, if we consider *Gfap*-expressing and *Slc8a2*-expressing EGCs separately), and all need to be stimulated in a coordinated way to ensure enteric neuronal diversity in well-balanced proportions. Most surprisingly, one of these progenitors seems not to derive from NCCs as virtually all ENS-related cells normally do.

### RET is dispensable for GDNF-induced neurogenesis in aganglionic distal colon

GDNF-RET signaling is critical for normal ENS formation from vagal NCCs ^55, 56^, explaining why *RET* variants are the main genetic risk factor for HSCR ^57–59^. However, modeling this association in mice has proved difficult. The impact of *Ret* deficiency on the formation of the murine ENS is generally either benign at heterozygous state ^60^ or too severe at homozygous state (total intestinal aganglionosis) ^56^. Distal colon aganglionosis mimicking short-segment HSCR can be induced by treating hypomorphic *Ret^9/-^* mice with mycophenolate mofetil ^61^, but general toxicity associated with this compound prevented us from evaluating GDNF effect in this context ^20^. Here, we used a pharmacological approach to circumvent these problems, demonstrating that RET activity is dispensable for mediating the neurogenic effect of GDNF in the aganglionic colon of *Holstein* mice. This result is also consistent with our scRNA-seq data showing that *Ret* expression is mostly restricted to a subset of induced neurons only. In marked contrast, *Ncam1* is robustly expressed in virtually all ENS progenitors and induced neurons, and pharmacological inhibition of the NCAM1-FAK signaling axis significantly impaired the neurogenic effect of GDNF.

NCAM1 was already known to influence ENS gangliogenesis before birth ^62^, and to mediate GDNF effects in the Schwann cell lineage after birth ^26^. Our new mouse data now suggest that NCAM1 also mediates GDNF effects in postnatal EGCs as well, at least in an aganglionosis context. There is also cumulative circumstantial evidence that NCAM1 can similarly mediate GDNF signaling in human tissues. In particular, NCAM1 (also known as CD-56) has been recently reported as a reliable marker for FACS-mediated recovery of ENS cells from human gastrointestinal tissues ^63^, including aganglionic samples obtained from children with HSCR ^47^. ScRNA-seq data generated from these NCAM1+ cells revealed the presence of both SCPs (referred to as Schwann-like enteric glia) and EGCs (referred to as canonical enteric glia) in aganglionic samples ^47^. These observations are also supported by our own immunofluorescence analyses of human aganglionic colon samples, which show robust NCAM1 protein expression both within (presumably in SCPs) and outside (presumably in EGCs) βIII-Tubulin+ extrinsic nerves (Fig.11).

**Figure 11.**
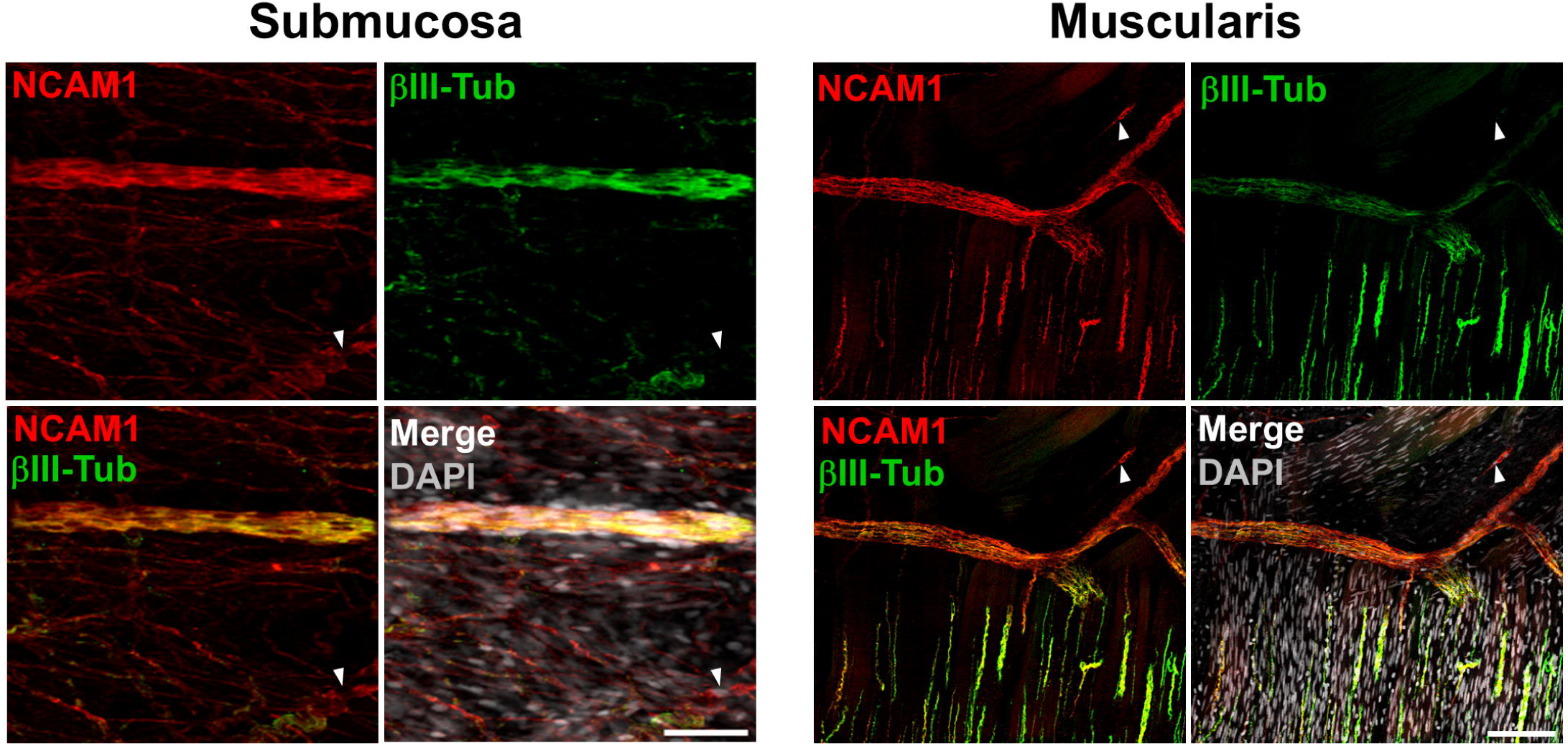
NCAM1 is robustly expressed in human samples of aganglionic colon. Immunofluorescence analysis reveals NCAM1 localization both within and outside βIII-Tubulin+ extrinsic nerves of the submucosa and muscularis tissue layers. Scale bar, 50 μm (submucosa) and 100 μm (muscularis).

The reason why RET is present in a subset of GDNF-induced neurons is unclear. RET is normally present in a subset of adult enteric neurons as well, but a recent study based on two distinct conditional mutant mouse lines (*Vip^Cre/+^*;*Ret^Flox/Flox^* and tamoxifen-inducible *Ret^CreER/Flox-GFP^*) suggests that it is not critical in these cells for ileal and colonic neuron survival or control of gastrointestinal motility ^64^. Yet, another study previously reported that RET inactivation in *Dhh*-expressing SCPs can reduce the number of neurons specifically in the distal colon of adult mice ^10^. Given that spontaneous SCP-derived neurogenesis is not impaired (being even more robust than normal) in the colon of *Ret^51^*^(C618F)*/−*^ mutant pups at P5 ^13^, this suggests that RET could be necessary for long-term survival of some SCP-derived neurons once induced.

Combined with our previous and current data, these observations are nonetheless reassuring about the global applicability of GDNF-based therapy, even in presence of lower RET activity. Indeed, even if RET is necessary for the long-term survival of some newly induced neurons or for mediating the presumed beneficial effects of GDNF in non-ENS cells (*e.g*., in epithelial cells ^64, 65^), it is noteworthy that HSCR-associated *RET* gene variants are generally predicted to reduce, but not abrogate, RET protein levels/activity ^58, 59^. In such cases with reduced RET levels/activity, GDNF-based therapy would still be highly valuable since GDNF enemas can trigger a feedback loop that increases protein levels of both RET and endogenous GDNF in the colon wall ^20^. This increase in endogenous GDNF is also interesting in relation to the persistence of neurogenesis after the end of GDNF treatment. Whether this is the only reason, or whether it implicates another signalling pathway (or a combination of the two) is certainly worthy of investigation in future research.

### The aganglionic distal colon is rich in ENS progenitors that can be directly converted into neurons by GDNF

This study provides additional evidence that aganglionosis does not mean absence of ENS progenitors ^66, 67^. Both mouse (this study) and human ^47^ scRNA-seq data uncovered two main progenitor subtypes that have the transcriptional signature of neural crest-derived SCPs or EGCs, as we further validated using several genetic cell lineage tracing tools. Yet, determining the exact relative contribution of each of these progenitor subtypes to the pool of GDNF-induced neurons is almost impossible to determine with currently available tools. The perfect models do not exist. For instance, we previously reported that ∼10% of SCPs from aganglionic colon are not fluorescently labeled when *R26R-YFP* reporter allele expression depends on the *Dhh-Cre* driver ^20^. Although efficacy of the GFAP-CreERT2 driver for depleting GFAP+ EGCs has been previously demonstrated during the early postnatal period ^68^, its tamoxifen-inducible nature could nonetheless also potentially underestimate the neuronal contribution of these EGCs beyond the tamoxifen administration window. However, our exploratory experiments suggest that this is not the case: extending tamoxifen administration until P18 (at 2-day intervals after P8; necessary to reduce deadly toxicity of chronic tamoxifen administration) resulted in a similar contribution of 33% of GDNF-induced neurons. On the other hand, the contribution of *Slc18a2*-expressing EGCs is likely overestimated with the *Slc18a2-Cre* driver, as this transgene is known to have intrinsic activity in various neuron subtypes in the central nervous system ^69–72^ and our scRNA-seq data show *Slc18a2* gene expression in the neuronal cluster EN #3. Moreover, the portrait is greatly complicated by the SCP-EGC lineage connection that we have discovered. In the end, the best estimate that we can make is based on the broader *Wnt1-Cre2* driver, showing that ∼75% of GDNF-induced neurons are collectively derived from NCCs.

This observation that not all GDNF-induced myenteric neurons are fluorescently labeled when using the pan-NCC driver *Wnt1-Cre2* was surprising. Indeed, in our hands, the *Wnt1-Cre2* driver normally turns on the *R26R-YFP* reporter allele in virtually all enteric neurons from the colon of WT-like control mice, and this pattern is even conserved in the “normoganglionic” proximal colon of *Holstein* mutant mice bearing the same tools and genetic background (FVB strain). Therefore, these data suggest that the non-NCC-derived ENS progenitor subtype that generates ∼25% of GDNF-induced neurons is uniquely present and/or responsive to GDNF in the diseased colon segment. The fact that this non-NCC-derived progenitor subtype differentiates preferentially into neurons (with increased capacity for nitrergic differentiation) rather than glial cells is also very intriguing. The quest to identify this unconventional progenitor cell population promises to be long and complex. Four alternative sources of ENS progenitors unrelated to the neural crest lineage have already been proposed over the years, including ventral neural tube ^73^, endodermal ^74^, mesodermal ^75^ and placodal ^76^ cell types. All of them were reported to be solicited during normal ENS development or aging. However, as mentioned above, our cell lineage tracing data with the *Wnt1-Cre2* driver do not support such conclusion for the ENS-containing colon of both WT-like and *Holstein* mouse pups before weaning, rather indicating full contribution by NCCs. That being said, such discrepancy might be due to diverse reasons related to experimental conditions/tools/settings, and these previously suggested unconventional progenitors could nonetheless represent legitimate candidates in the aganglionic colon soon after birth. Of note, inspiration could also come from a recent preprint reporting the spontaneous generation of enteric neurons in human intestinal organoids without the need to add NCCs ^77^. Clearly, there will be a lot of potential candidates (both known and not yet known) to experimentally test in future work using a framework similar to the current study.

Another intriguing discovery that was made during these studies is that direct transdifferentiation is the main mode of GDNF-induced neurogenesis in the aganglionic distal colon regardless of ENS progenitor subtypes and time point relative to treatment period. This contrasts with prior studies in WT mice in which systemic subcutaneous administration of GDNF between P0 to P30 expanded the submucosal neuron pool (but not the myenteric pool) by increasing the proliferation rate of submucosal neuronal precursors ^78^, a finding also corroborated by the study of *Gdnf^+/-^* mice ^60^. Yet, we also observed an increase of such “indirect” differentiation via a dividing precursor over time, thereby most likely preventing GDNF to rapidly exhaust the pool of ENS progenitors. Such delayed response suggests that the pool of ENS progenitors is somehow protected by a quorum sensing-like mechanism that controls cellular density at tissue level, as observed in the immune system ^79^. Regardless of exact mechanism, the resulting two-step response is highly desirable from a therapeutic standpoint, as it allows to combine both rapid and long-lasting effects of GDNF treatment.

Finally, our discovery that multiple progenitor subtypes are required to obtain a properly balanced ENS also has important implications for all cell transplantation-based approaches aimed at replacing missing neurons. This means that the subtype(s) of ENS stem/progenitor cells to transplant should be selected with care, as a function of the neuronal subtype to replace.

## MATERIALS AND METHODS

### Sex as biological variable

Both female and male individuals were used in all experiments involving mouse or human samples, and similar findings are reported for both sexes.

### Mice

*Holstein* (*Tg[Sox3-GFP,Tyr]HolNpln*) and *G4-RFP* (*Gata4p[5kb]-RFP*) transgenic lines were as previously described in our prior work ^29, 34^. *Dhh-Cre* (*FVB[Cg]-Tg[Dhh-cre]1Mejr/J*; JAX strain #012929), *GFAP-CreERT2* (*B6.Cg-Tg[GFAP-cre/ERT2]505Fmv/J*; JAX strain #012849) and *Wnt1-Cre2* (*B6.Cg-E2f1^Tg(Wnt1-cre)2Sor^/J*; JAX strain #022501) mouse lines were purchased from the Jackson Laboratory (Bar Harbor, MM, USA). The *Slc18a2-Cre* (*Tg[Slc18a2-Cre]OZ14Gsat/Mmucd*; MMRRC strain #034814-UCD) line was obtained from the Mutant Mouse Resource and Research Centers (MMRRC) at University of California at Davis, an NIH-funded strain repository, and was donated to the MMRRC by Nathaniel Heintz (The Rockefeller University, GENSAT) and Charles Gerfen (National Institutes of Health, National Institute of Mental Health) ^80^. The Cre reporter line *Rosa26^FloxSTOP-YFP^* (*B6.129X1-Gt[ROSA]26Sor^tm1(EYFP)Cos^/J*) was directly provided by Dr. Frank Costantini from Columbia University. All mice used in this study were maintained on the FVB/N genetic background, after at least 5 generations of backcrossing if acquired on a different background. All mice were housed and bred in individually ventilated cages within the conventional animal facility at the Université du Québec à Montréal, under a 12-hour light/dark cycle (7 AM to 7 PM) and with *ad libitum* access to Charles River Rodent Diet #5075 (Cargill Animal Nutrition). Genotyping was either made by visual inspection of coat color (for *Holstein* and *G4-RFP*) or standard PCR (for all Cre driver and reporter lines) using primers listed in Table S1. Euthanasia was performed using either CO₂ inhalation or decapitation (for pups aged of 10 days or less), both following isoflurane anesthesia.

### Administration of substances

GDNF enemas (10µl at 1mg/ml in PBS) were administered once daily between P4-P8, using a 24-gauge gavage needle (Fine Science Tools, Canada) connected to a micropipette, as previously described ^20, 21^. Briefly, gavage needle tip was gently inserted into the rectum just beyond the anus, and enema was carefully injected over a few seconds. Two batches of human recombinant GDNF were used, both standardly produced in bacteria. Research-grade GDNF purchased from PeproTech (cat. #450-10, Cranbury, NJ) was used for single-cell RNA-seq experiments, while clinical-grade GDNF kindly provided by Vivifi Biotech (London, United-Kingdom) was used for all other experiments. When required, tamoxifen (20µl at 4mg/ml in corn oil; Sigma Cat.#10540-29-1) and/or EdU (10 µl at 10mM in PBS; Thermo Fisher Cat.# C10337) were administered via daily intraperitoneal injections in parallel to GDNF enemas, but starting a day earlier (P3-P8). Phospho-FAK[Y397] and phospho-RET[Y1062] inhibitors, PF-562271 (MedChemExpress Cat.# HY-10459) and Pralsetinib/BLU-667 (MedChemExpress Cat.# HY-112301), respectively, were similarly administered via daily intraperitoneal injections using same 30mg/kg concentration (in a vehicle solution of 2% DMSO; 25% PEG 300; 5% Tween 80; 68% Saline), but in this case only during GDNF treatment period (P4-P8).

### Human Tissues

Human colon samples were obtained from three patients (2 males, 1 female) diagnosed with short-segment HSCR who underwent surgical resection of their aganglionic colon segment. These patients were recruited at the Centre hospitalier universitaire Sainte-Justine and their ages ranged between 3-6 months at time of surgery. Informed consent for collection and use of these tissues was obtained from all parents or legal guardian. Following surgical resection, full-thickness colon samples of aganglionic segment were immediately placed in ice-cold, oxygenated Krebs solution and promptly transported to UQAM research laboratory for further processing.

### Single-cell RNA-seq

Samples of the distal third of the colon (including part of transition zone) were dissected from P10 *Hol^Tg/Tg^;G4-RFP* double-transgenic pups that were previously administered GDNF enemas or not between P4-P8. For each condition, colon samples were collected from 6 to 8 pups and pooled on day of dissection. Once pooled, each piece of distal colon was cut opened along longitudinal axis, washed extensively in ice-cold PBS, and mechanically dissociated during 5 minutes using a scalpel blade in a small volume of pre-warmed (37°C) EMEM (Eagle’s Minimum Essential Medium) containing collagenase (0.4 mg/ml; Sigma-Aldrich C2674), dispase II (1.3 mg/ml; Life Technologies 17105-041), and DNAse I (0.5 mg/ml; Sigma-Aldrich DN25). Resulting grossly dissociated tissue preparation was transferred in an Eppendorf tube, into which the above dissociation medium was added to reach ∼1 ml total volume, and the tube was then placed in an agitating heating block at 37°C for 30 minutes at 300rpm. Final tissue preparation was filtered through a 40µm cell strainer and centrifuged at 1200 rpm for 5 minutes at 4°C. Pelleted cells were resuspended in 500 µl of sterile PBS also containing 0.04% non-acetylated BSA, stained with CellTracker™ Green CMFDA Dye (Invitrogen, Cat.#C2925) diluted at 1/10000, and separated in a BD FACSJazz cell sorter (BD Biosciences) to recover viable CMFDA+ RFP+ cells, under most permissive Yield mode parameters (Yield Mask = 32 and Purity Mask = 0). Single-cell RNA-seq libraries were prepared following the guidelines of the 10x Genomics protocol. Brefly, single-cell Gel Bead-In-Emulsions (GEMs) were generated by combining suspended cells, barcoded gel beads, and master mix onto the Chromium Next GEM Chip G, which was then processed on the Chromium iX controller Instrument. After breaking the GEMs, barcoded cDNAs were purified and amplified according to desired cell recovery target. Subsequently, the cDNA underwent fragmentation, A-tailing, and ligation with adaptors. For sample indexing, additional PCR cycles were conducted based on cDNA input quantified during QC step. Resulting libraries were finally subjected to 100bp paired-end sequencing on an Illumina NovaSeq 6000 sequencer.

### Single-cell RNA-seq data analysis

Single-cell RNA-seq data were analyzed using Trailmaker (Parse Biosciences). After alignment to the reference genome (mm10) and transcript quantification, quality filtering was applied to exclude low-quality cells. The initial dataset contained 10,018 to 9,477 barcodes, depending on the experimental conditions, with a total of 22,142 to 22,778 detected genes. A mitochondrial content filter was applied, setting a maximum threshold of 10% mitochondrial reads with a bin step of 0.30. After filtering, the number of retained cells decreased from 10,018 to 3,997 in the untreated *Hol^Tg/Tg^* group (-60.1%) and from 9,477 to 7,091 in the GDNF-treated *Hol^Tg/Tg^* group (-25.2%). Data normalization and batch effect correction were performed using the integrated Trailmaker tools. Dimensionality reduction was conducted via principal component analysis (PCA), and cell clustering was performed with the Leiden algorithm. Cell populations were visualized using uniform manifold approximation and projection (UMAP). Cluster annotation was performed by comparing differentially expressed marker genes with reference databases (PanglaoDB, CellMarker). Differential gene expression analysis was carried out using the Wilcoxon test implemented in Trailmaker. Cell trajectory reconstruction was conducted using Trailmaker’s pseudotime tools.

### Tissue processing, labelling and imaging

Mouse colon tissues to be immunostained were cut longitudinally along the mesentery, washed in PBS, pinned onto Sylgard-coated (Thermo Fisher Scientific Cat. #50822180) petri dishes, fixed with 4% PFA at 4°C overnight, and finally further microdissected to separate muscle and mucosa layers. Whole muscle strips of the most distal quarter of the colon were then used for immunofluorescence staining, which was performed as previously described ^20^. For human colon tissues, immunofluorescence staining was performed on full-thickness aganglionic samples using a previously described protocol for 3-dimensional imaging ^81^. All antibodies used for immunostaining are listed in Table S2. EdU detection in mouse samples was performed using Invitrogen Click-iT EdU Imaging Kit (Thermo Fisher Scientific Cat. #C10337), in accordance with manufacturer’s instructions. All images were acquired on a Nikon A1 confocal microscope, using either Plan Fluor 20x/0.75 MImm or Plan Apo λ 60x/1.40 objectives (Nikon A1R, Melville, NY). For each biological replicate, 3 to 11 representative z-stack images of myenteric plexus were acquired, and then analyzed using the ImageJ software (National Institutes of Health, Bethesda, MD).

### Western blot analysis

Remaining part of colon samples not used for immunofluorescence (*i.e.*, proximal and mid segments) were homogenized in RIPA buffer containing protease and phosphatase inhibitors and centrifuged (14000rpm for 15min at 4C) to sediment debris. Protein concentration in supernatant was determined using DC protein assay (Bio-Rad Kit I #5000111) according to manufacturer’s guidelines. For western blot analysis, 50 µg of total protein extracts were separated on a 10% SDS-PAGE and then transferred onto an Immun-Blot PVDF membrane (Bio-Rad Cat. #1620177). Immunoreactive bands were revealed using Immobilon Western Chemiluminescent HRP Substrate (MilliporeSigma Cat. #WBKLS0050) and visualized using the Fusion FX imaging system (Vilber, Marne-la-Vallée, France). All antibodies used are detailed in Table S2.

### Data statistics

Statistical analyses were performed using GraphPad Prism v9.5.1. As indicated in Figure legends, either Student’s *t*-tests or one-way ANOVA with post-hoc Tukey’s test were used to calculate statistical significance, with *P* values < 0.05 considered as indicative of significance. All experiments included a minimum of 3 biological replicates.

### Study approval

All experiments involving mice were approved by the Université du Québec à Montréal ethics committee (CIPA #959) and conducted in accordance with the Canadian Council on Animal Care (CCAC) guidelines. Use of human samples in our experiments strictly adhered to guidelines and approval of human research ethics committee of the Centre Hospitalier Universitaire Sainte-Justine (CER protocol #4172).

### Data availability

All relevant data are included in the article. Raw files of single-cell transcript profiling can be accessed on the Gene Expression Omnibus database (series GSE313116). For reagent requests, please contact RS or NP.

## AUTHOR CONTRIBUTIONS

Conceived and Supervised the study: RS and NP. Designed the experiments: AG, RS and NP. Contributed study materials: AA and CF. Performed the experiments: AG, MAL, NL and RS. Analyzed the data: AG, RS and NP. Wrote the paper: AG (original draft), RS (original draft and review), and NP (review and editing). All authors read and approved the final manuscript.

## ACKNOWLEDGEMENTS

The authors thank Gregoire Bonnamour from the Cellular analyses and Imaging core (CERMO-FC, UQAM) for assistance with flow cytometry and confocal imaging, and Farzaneh Ramdani from the Bioinformatics core (CERMO-FC, UQAM) for help with scRNA-seq data processing.

