## Supplementary material for "GDNF regenerates the missing enteric nervous system of Hirschsprung mice via non-canonical signaling in diverse subtypes of tissue-resident progenitors": Table S1

*Gary et al.*

**SUPPLEMENTAL MATERIAL**

**Table S1. Specific primers used for genotyping.**

**Table S2. List of primary antibodies used in this study.**

**Table 1. Specific primers used for genotyping.**

| **ID** | **Sense primer** | **Antisense primer** |
| --- | --- | --- |
| *Cre* | 5’-GATGAGGTTCGCAAGAACCTGATG | 5’-AACAGCATTGCTGTCACTTGGTCG |
| *R26R-YFP* | 5’-CCCAAAGTCGCTCTGAGTTGTTATC | YFP: 5’-TGCGCCCTACAGATCCCTTAATTAA  WT: 5’-CCAGATGACTACCTATCCTCCCA |
| *Holstein* | 5’-GTGGTGGACCTAACCTTACAAGGA | WT: 5’-CAGGGCTAAGTCTTGGCTTACTTG  Mut: 5’-CACAGCTTGCTGTATCAGAGCCAT |

**Table 2. List of primary antibodies used in this study.**

| **Primary Antibody** | **Source** | **Catalog number** | **Species** | **RRID** | **Dilution** |
| --- | --- | --- | --- | --- | --- |
| **Immunofluorescence** | | | | | |
| Anti-βIII-Tubulin | Abcam | ab78078 | Mouse | AB_2256751 | 1:500 |
| Anti-HuC/D | Molecular Probes | A-21271 | Mouse | AB_221448 | 1:500 |
| Anti-SOX10 | In house (MediMabs) | N/A | Rat | N/A | 1:500 |
| Anti-SOX10 | R&D Systems | AF2864 | Goat | AB_442208 | 1:500 |
| Anti-ChAT | Millipore | AB144P | Goat | AB_2079751 | 1:250 |
| Anti-GFP | Abcam | ab290 | Rabbit | AB_303395 | 1:500 |
| Anti-GFP | Abcam | ab6673 | Goat | AB_305643 | 1:500 |
| Anti-NCAM | Millipore | AB5032 | Rabbit | AB_2291692  AB_11213653 | 1:500 |
| Anti-NOS1 | Millipore | AB5380 | Rabbit | AB_91824 | 1:500 |
| Anti-PHOX2B | Novus | AF4940 | Goat | AB_2861427 | 1:250 |
| Anti-pFAK[Y397] | Abcam | ab81298 | Rabbit | AB_1640500 | 1:250 |
| Anti-GFAP | Novus | NB100-53809 | Goat | AB_829022 | 1:500 |
| Anti-Caspase-3 | Abcam | ab13847 | Rabbit | AB_443014 | 1:500 |
| **Western Blot** | | | | | |
| Anti-RET | R&D Systems | AF482 | Goat | AB_2301030 | 1:1000 |
| Anti-pRET[Y1062] | Abcam | ab51103 | Rabbit | AB_870738 | 1:200 |
| Anti-FAK | Millipore | 05-537 | Mouse | AB_2173817 | 1:1000 |
| Anti-pFAK[Y397] | Abcam | ab81298 | Rabbit | AB_1640500 | 1:1000 |
| Anti-GAPDH | Santa Cruz | sc-32233 | Mouse | AB_627679 | 1:5000 |

RRID: Research Resource Identifiers. N/A: not applicable
